# Heritability and relationship of oxytocin receptor gene variants with social behavior and central oxytocin in colony-reared adult female rhesus macaques

**DOI:** 10.1101/813287

**Authors:** Desirée De Leon, Hasse Walum, Kai M. McCormack, Jamie L. LaPrairie, Stephen G. Lindell, Ashley Firesryche, Zach Johnson, Mark E. Wilson, Christina S. Barr, Mar M. Sanchez, Larry J. Young

## Abstract

The genetic contributions to sociality are an important research focus for understanding individual variation in social function and risk of social deficits in neurodevelopmental disorders (e.g. autism). The neuropeptide oxytocin (OXT) and its receptor, OXTR, influence social behavior across species. In humans and animals, common variants within the OXTR gene (*OXTR*) have been associated with varying socio-behavioral traits. However, the reported magnitude of influence of individual variants on complex behavior has been inconsistent. Compared to human studies, non-human primate (NHP) studies in controlled environments have the potential to result in robust effects detectable in relatively small samples. Here we estimate heritability of social behavior and central OXT concentrations in 214 socially-housed adult female rhesus macaques, a species sharing high similarity with humans in genetics, physiology, brain and social complexity. We present a bioinformatically-informed approach for identifying single nucleotide polymorphisms (SNPs) with likely biological relevance. We tested 13 common SNPs in regulatory and coding regions of *OXTR* for associations with behavior (pro-social, anxiety-like, and aggressive) and OXT concentration in cerebrospinal fluid (CSF). We found moderate rates of heritability for both social behavior and CSF OXT concentrations. No tested SNPs showed significant associations with behaviors or CSF in this sample. Associations between OXT CSF and social behavior were not significant either. SNP effect sizes were generally comparable to those reported in human studies of complex traits. While environmental control and a socio-biological similarity with humans is an advantage of rhesus models for detecting smaller genetic effects, it is insufficient to obviate large sample sizes necessary for appropriate statistical power.

## Introduction

Oxytocin is an evolutionarily conserved neuropeptide involved in orchestrating reproductive, maternal, and social behaviors across species [1]. Individual variability in these traits is attributable to genetic variation, apart from the role of experience. As such, variants such as single nucleotide polymorphisms (SNPs) within the OXT receptor gene (*OXTR*) may contribute to the diversity of complex social repertoires. Genetic association studies suggest that variation in *OXTR* may explain part of the variance in social phenotypes including pair-bonding behavior, social recognition, prosocial temperament, and sensitivity to social stressors [1–3]. Multiple *OXTR* SNPs have also been linked to social impairments in individuals with autism [4].

Despite reports linking human *OXTR* variants to social phenotypes, other studies, including a meta-analysis [5], have failed to find effects of consistent magnitude and direction of these variants on social domains [see 6 for review]. Inconsistent results may be related to biases which disproportionately suppress non-significant results from publication (e.g. publication bias, selective reporting), and inflated effect sizes due to low statistical power [7].

The use of animal models, however, has allowed the underlying mechanisms for SNP-behavior associations to be probed in the brain, including assessing the functional effects of differential *OXTR* gene expression caused by genetic variants in brain tissue. For example, in the prairie vole rodent model, a single *Oxtr* SNP predicted more than 70% of the variability in expression in the nucleus accumbens, a region critical for social reward [8]. This suggests that individual variants have the potential to profoundly impact brain phenotypes which can mediate downstream behavior differences.

Non-human primates (NHPs), such as the rhesus macaque, have been widely used to study social behavior. Both rhesus and humans display rich behavioral repertoires, complex social hierarchies, and are very similar in their physiology and neuroanatomy. Additionally, rhesus have frequently been used to investigate the role of OXT on sociality [9], and as such, an understanding of the genetic contributions to the oxytocinergic system in this NHP model is valuable for translation to humans.

While human genetic association studies necessitate extremely large samples to detect small effects reliably among the multitude of factors influencing complex traits, it is possible that NHP studies conducted in experimentally controlled, semi-naturalistic settings with known pedigrees can detect larger effects due to the reduction in environmental noise (e.g. standard diets and perinatal environments). The goal of the present study was to use an informed approach to examine the associations of novel rhesus *OXTR* (*rhOXTR*) SNPs with social behavior (pro-social, anxiety-like, and aggressive) and OXT in cerebrospinal fluid (CSF) in 214 adult female rhesus macaques housed in large social groups. We also report estimates of heritability and examine the effects of OXT CSF on each behavior.

## Materials and methods

### Subjects and housing

Subjects were 214 adult female rhesus monkeys (4-24 years in age; median: 7, interquartile range: 6-11) of known pedigrees living in complex social groups at the Yerkes National Primate Research Center (YNPRC) Field Station (Lawrenceville, GA) as described in [10]. Subjects came from five social groups consisting of 28-94 adult females, their kin, and 2-10 adult males. Animals were selected from matrilines across all social status ranks (high, medium and low) and with varying degrees of relatedness for heritability analyses. Animals were housed in outdoor enclosures (3/4 to 1 acre areas) with access to climate-controlled indoor facilities and provided a standard commercial low-fat, high-fiber diet (Purina Mills International, LabDiets, St. Louis, MO) and water *ad libitum*, supplemented with seasonal fruits or vegetables twice per day. All procedures complied with the Animal Welfare Act and U.S. DHHS “Guide for the Care and Use of Laboratory Animals” and were approved by the Emory IACUC.

### Behavioral data collection

Focal observations were collected in real-time following published protocols [10,11] based on established ethograms (Altmann, 1962). An average behavioral observation of 80 min/animal was collected during the mating season to reduce seasonal variability in behavior. We focused on representative behaviors: pro-social (percent of time spent in proximity to other adult females), anxiety-like (frequency of self-scratches) [12], and aggressive (non-contact, subject-initiated) behaviors (Table 1).

**Table 1.** Specific behaviors analyzed for this study.

| Variable | Definition | Data Collected |
| --- | --- | --- |
| ***Pro-social Behavior*** |  |  |
| Proximity to other adult females<sup>a</sup> | Subject is within 1 foot of one or more other adult females (including physical contact) | Duration |
| ***Aggressive Behavior*** |  |  |
| Non-contact aggression | Counts of agonistic chases, open-mouth threats, barks, and lunges of/by another animal | Frequency |
| ***Anxiety-like / Solitary Behavior*** |  |  |
| Anxiety/self-directed | Counts of self-scratches<sup>b</sup> | Frequency |

<sup>a</sup> Given the matrilineal structure of rhesus troops [19], proximity to adult females (as opposed to other members in the group) comprise the most frequent and relevant social interaction observed
<sup>b</sup> In order for the same frequency behavior to be scored a second time, at least 3 seconds had to have passed from the first instance of the behavior

### CSF and blood samples

Animals were habituated to experimental procedures to facilitate blood and CSF collection using methods minimizing arousal [10]. Each subject was accessed once, shortly after sunrise and during mating season, but never on the same day as the behavioral data collection. Animals were anesthetized with Telazol (3 mg/kg, IM). CSF samples were collected in a subset of the subjects (n=166) to examine central OXT concentrations. CSF samples were collected (2 ml/subject) from the *cisterna magna* by gravity through a 22 G needle and placed immediately on dry ice. Two 3 ml blood samples were collected in EDTA tubes for DNA extraction and immediately placed on ice. Samples were stored at -80°C until time of processing. CSF samples were processed by the YNPRC Biomarkers Core Laboratory using commercially available ELISA kits produced by Assay Designs (Ann Arbor, MI), following manufacturer’s recommendations. Sensitivity of the OXT assay was 15.6 pg/ml and the inter- and intra-assay CVs were 7.48% and 10.2%, respectively.

### Covariates

Social rank and age (and no other variables) were included as covariates for all models. OXT concentrations were assayed in three batches; observations were mean-centered per batch to control for inter-batch variation.

### SNP selection

SNP discovery efforts were completed in a separate group of rhesus and focused on 5’ regulatory and coding regions of *rhOXTR.* Within these regions, candidate SNPs were identified in positions analogous to those in humans that have been cited as being involved in *OXTR* expression [13] or suggested to predict individual differences in behavior and disease vulnerability. While SNPs are not conserved across species, comparably located SNPs in macaques may confer functionally similar effects and therefore provide translational potential. Additional SNPs in highly conserved areas within the region of interest were also included as candidates. Candidates were narrowed down based on their presence with sufficient minor allele frequency in that separate group of macaques, comprising representatives from multiple genetically-distinct populations. This resulted in 13 loci to be genotyped for this study’s sample (Fig 1).

**Fig 1.**
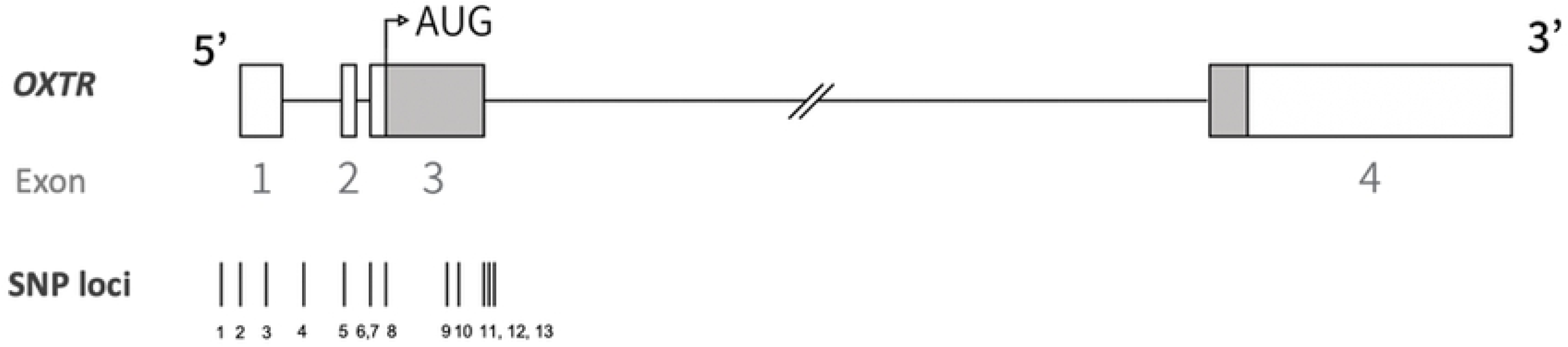
Schematic of OXTR SNPs probed in this study. Untranslated portions of the exons are indicated by the white bars. Genomic coordinates correspond to chromosome 3 of the MacaM reference genome (SNP 1) 140481032, (SNP 2) 140497572, (SNP 3) 140497358, (SNP 4) 140497003, (SNP 5) 140496605, (SNP 6) 140496348, (SNP 7) 140496344, (SNP 8) 140496196, (SNP 9) 140495645, (SNP 10) 140495542, (SNP 11) 140495303, (SNP 12) 140495244 G/A, (SNP 13) 140495203.

### Genotyping

Three different genotyping techniques were used. Archived next-generation sequencing data targeting *OXTR* exons was available for loci 4-13. Libraries were generated using the Illumina NexteraXT DNA kit and sequenced on an Illumina HiSeq1000. The remaining markers were genotyped using TaqMan or Sanger Sequencing. Cycle sequencing was performed using the Big Dye Terminator, version 3.1, reaction in 96-well optical plates (Applied Biosystems, Foster City, California). Variants were detected by visualization of electropherograms generated by ABI Sequencing Analysis software. Relevant assay and primer information are in Supplemental Table S1.

### Statistical analysis

All statistical analyses were performed in R version 3.4.1 (R Project for Statistical Computing, Vienna, Austria). Associations of SNPs with behavioral and CSF measures, as well as heritability, were examined using the “animal model” with the “MCMCglmm” package [14]. Narrow-sense heritability was calculated as the proportion of variance explained by additive genetic factors (via the pedigree) out of all phenotypic variance. SNPs and covariates were included as fixed effects, and relatedness was accounted for by including pedigree as a random effect. To account for differences in total time observed, observation length was included as an offset for models including frequency behaviors (i.e. raw counts). The prior distribution for additive genetic variance (σ^2^_A_) was defined by commonly used non-informative parameters for the inverse-gamma distribution, IG(0.001, 0.001). Models were run with a minimum of 5 million iterations, a burn-in of 5,000, and thinning interval of 1,000 until adequate mixing was ensured.

A Gaussian distribution was specified for pro-social behavior and log-transformed OXT CSF measures. For anxiety-like behavior, a Poisson specification was used. Heritability was not estimable for aggression, even after trying multiple distributions and stronger priors on σ^2^ _A_ This was most likely due to a combination of low occurrence of this behavior (i.e. low information) and a small sample size [15]. Because controlling for relatedness was essential for all models, this outcome was excluded from subsequent analyses.

In post-hoc analyses, a likelihood-ratio test was used to compare whether inclusion of all SNPs (versus none) significantly improved model fit. To do so, frequentist versions of each model were run using R packages “pedigreemm” and “regress” to extract maximum log-likelihood values.

## Results

### Heritability estimates

We report moderate heritability for pro-social behavior (h^2^ = 0.312), anxiety-like behavior (h^2^ = 0.283), and OXT CSF (h^2^ = 0.183), though all 95% credible intervals were wide (Table 2).

**Table 2.** Heritability and SNP estimates.

| Narrow sense heritability ($h^2$) | Proximity to Adult Females (Pro-social) | | | Self-Scratches (Anxiety-like) | | | OXT in CSF | | |
| --- | --- | --- | --- | --- | --- | --- | --- | --- | --- |
|  | Point Estimate | Lower 95% CI | Upper 95% CI | Point Estimate | Lower 95% CI | Upper 95% CI | Point Estimate | Lower 95% CI | Upper 95% CI |
| Without fixed effects | 0.312 | 0.046 | 0.591 | 0.213 | $4.98 \times 10^{-4}$ | 0.670 | 0.183 | 0.002 | 0.478 |
| With fixed effects | 0.306 | 0.038 | 0.587 | 0.297 | $3.95 \times 10^{-4}$ | 0.679 | 0.193 | $7.93 \times 10^{-4}$ | 0.500 |
| SNPs | $\beta$ | Lower 95% CI | Upper 95% CI | $\beta$ | Lower 95% CI | Upper 95% CI | $\beta$ | Lower 95% CI | Upper 95% CI |
| 1) 140481032 G/A | -0.011 | -0.043 | 0.023 | 0.003 | -0.141 | 0.148 | 0.039 | -0.056 | 0.136 |
| 2) 140497572 G/C | 0.003 | -0.028 | 0.037 | 0.058 | -0.089 | 0.224 | 0.023 | -0.072 | 0.123 |
| 3) 140497358 G/A | 0.003 | -0.028 | 0.038 | 0.079 | -0.076 | 0.238 | 0.025 | -0.074 | 0.122 |
| 4) 140497003 G/T | -0.037 | -0.132 | 0.063 | -0.481 | -0.912 | 0.015 | 0.123 | -0.177 | 0.431 |
| 5) 140496605 T/C | -0.006 | -0.038 | 0.027 | 0.043 | -0.126 | 0.19 | 0.015 | -0.073 | 0.108 |
| 6) 140496348 A/G | -0.004 | -0.038 | 0.027 | 0.031 | -0.111 | 0.188 | 0.019 | -0.073 | 0.108 |
| 7) 140496344 A/C | -0.005 | -0.038 | 0.028 | 0.032 | -0.126 | 0.181 | 0.018 | -0.076 | 0.115 |
| 8) 140496196 G/T | 0.014 | -0.026 | 0.049 | 0.043 | -0.128 | 0.22 | -0.032 | -0.131 | 0.072 |
| 9) 140495645 C/G | -0.007 | -0.036 | 0.021 | 0.107 | -0.037 | 0.242 | 0.038 | -0.048 | 0.131 |
| 10) 140495542 A/T | -0.005 | -0.096 | 0.087 | 0.056 | -0.386 | 0.526 | 0.126 | -0.117 | 0.387 |
| 11) 140495303 C/T | 0.004 | -0.028 | 0.034 | -0.064 | -0.21 | 0.084 | 0.06 | -0.035 | 0.157 |
| 12) 140495244 G/A | -0.005 | -0.052 | 0.037 | -0.176 | -0.381 | 0.04 | 0.044 | -0.104 | 0.17 |
| 13) 140495203 T/G | -0.001 | -0.033 | 0.03 | -0.076 | -0.229 | 0.072 | 0.068 | -0.027 | 0.154 |
| | $\beta$ | Lower 95% CI | Upper 95% CI | $\beta$ | Lower 95% CI | Upper 95% CI | | | |
| OXT CSF vs Behavior | -0.024 | -0.080 | 0.032 | -0.273 | -0.543 | 0.006 | - | - | - |

### SNP associations

All 13 markers investigated (Fig 1) conformed to Hardy-Weinberg equilibrium, except two (SNPs 1 and 9, Table 3). No significant associations were detected between individual markers and behavioral outcomes or OXT CSF in our sample, as all uncorrected 95% credible intervals overlapped zero (Table 2). When all markers were included in the model, no significant improvement in model fit was observed, indicating that the cumulative effect of all SNPs still results in negligible effects on the investigated traits.

**Table 3.**
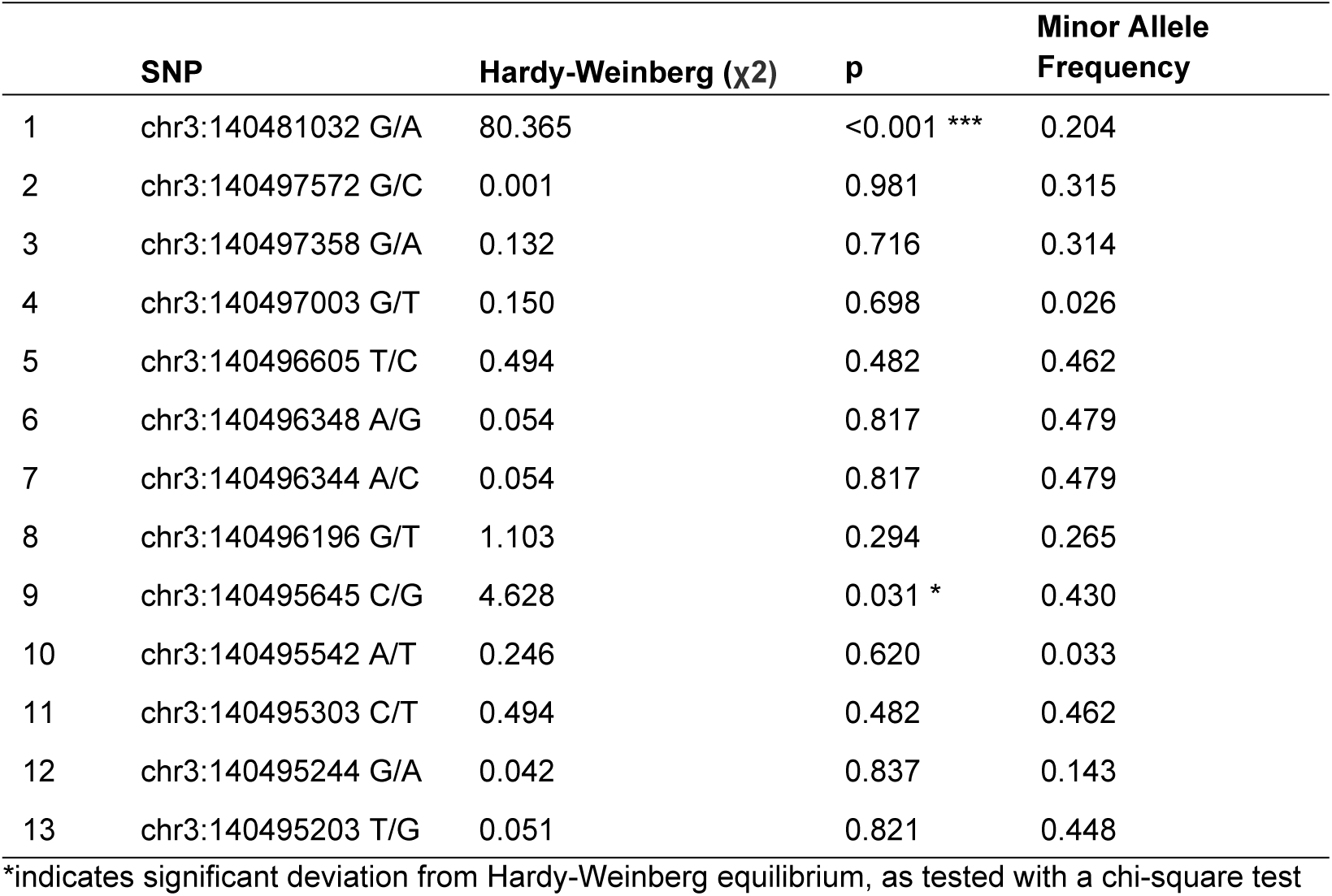
*OXTR* SNP Characteristics.

| | SNP | Hardy-Weinberg ($\chi^2$) | p | Minor Allele Frequency |
| --- | --- | --- | --- | --- |
| 1 | chr3:140481032 G/A | 80.365 | <0.001 *** | 0.204 |
| 2 | chr3:140497572 G/C | 0.001 | 0.981 | 0.315 |
| 3 | chr3:140497358 G/A | 0.132 | 0.716 | 0.314 |
| 4 | chr3:140497003 G/T | 0.150 | 0.698 | 0.026 |
| 5 | chr3:140496605 T/C | 0.494 | 0.482 | 0.462 |
| 6 | chr3:140496348 A/G | 0.054 | 0.817 | 0.479 |
| 7 | chr3:140496344 A/C | 0.054 | 0.817 | 0.479 |
| 8 | chr3:140496196 G/T | 1.103 | 0.294 | 0.265 |
| 9 | chr3:140495645 C/G | 4.628 | 0.031 * | 0.430 |
| 10 | chr3:140495542 A/T | 0.246 | 0.620 | 0.033 |
| 11 | chr3:140495303 C/T | 0.494 | 0.482 | 0.462 |
| 12 | chr3:140495244 G/A | 0.042 | 0.837 | 0.143 |
| 13 | chr3:140495203 T/G | 0.051 | 0.821 | 0.448 |

\*indicates significant deviation from Hardy-Weinberg equilibrium, as tested with a chi-square test

### Associations between OXT CSF and behavior

Finally, the relationship between OXT CSF and pro-social and anxiety-like behavior yielded no significant effects (Table 2).

## Discussion

Rhesus are a species with great translational value due to their behavioral and biological parallels to humans. Animals in this particular study also experienced complex social housing, and experimentally controlled environments (similar living, housing, and dietary conditions), attenuating the variability introduced by environmental factors that impact human studies. These benefits might suggest that rhesus behavioral genetic studies, which aim to detect subtle genetic signals among various sources of environmental noise, are exempt from the large samples sizes necessary in analogous studies conducted in humans. However, the use of the rhesus model did not produce effects large enough to be detected in this sample. Specifically, we report no significant associations between the 13 *rhOXTR* SNPs on pro-social or anxiety-like behavior, or on OXT concentrations in CSF. Further, OXT CSF was not associated with any behavioral outcomes.

Although we did not find significant effects of individual markers, our heritability analysis showed moderate additive genetic effects on all investigated traits, underscoring the existence of substantial genetic influence on these outcomes. This is unsurprising given extensive research in humans demonstrating that most complex traits are on average 50% heritable [16].

Regarding SNP associations, explanations for our null results include: (1) Despite our informed strategy for selecting markers, it is possible we identified *rhOXTR* variants that do not affect expression (nor subsequent downstream normative behavior). This is a limitation of candidate-gene approach, which narrows the breadth of testable markers. Whole-genome association methods would be needed to comprehensively identify any and all robust markers (while acknowledging the non-trivial challenges to statistical power this introduces). Alternatively (2), the contributions of individual SNPs across the genome, including these 13, may have small, incremental effects on polygenic, complex traits that studies like this one are too underpowered to detect. Decades of research in humans indicates the latter is more plausible [17]. As such, the experimental control afforded by NHP studies such as this one is not sufficient to generate effects robust enough to override the necessity for large samples required in human studies.

Our results are consistent with [18], who also investigated *rhOXTR* markers and social behavior in a similarly-sized sample of rhesus and likewise found no significant effects. We agree with their conclusions that behavioral genetic studies in NHPs likely face the same challenges as in humans: First, samples upwards of tens of thousands would be required to be adequately powered. Second, reported associations with complex traits resulting from small genetic studies are more likely to be false-positives and fail to replicate, which has been well-documented in human literature.

While our approach to select influential *rhOXTR* markers was not effective for our specific behavioral outcomes in the context of this sample size, it could be suitable for identifying endophenotypes such as brain expression and neural activity, which bear a closer biological relationship to the proximate consequences of genetic variation. Future genetic associations studies of social behavior in NHPs should parallel the human genetics field in shifting to extensive collaborative efforts, resulting in the possibility of acquiring large sample sizes, surpassing what would be feasible for individual research groups. Importantly, our results do not negate the wealth of OXT research demonstrating OXT’s role in regulating social behavior, nor do they refute the possibility that *OXTR* SNPs may have measurable effects in larger samples, or under different experimental conditions (e.g. stress challenges). Whole-genome approaches, when appropriate sample sizes can be attained, can address the relative influence of *OXTR* variants on shaping primate social behavior.

## Acknowledgments

We like to thank Anne Glenn, Jennifer Miles, Geary Smith, Jeff Fisher, Jennifer Whitley, Amy Henry, Krystle Ainsworth, Colin White, and Sneka Raveendran for their technical support as well as the Yerkes NPRC Biomarkers Core for performing bioassays. We would also like to thank Dr. Jeffrey Rogers for his helpful insights with the experimental design of this study.

## Supporting information

**Table S1. Assay and Primer information**

